# The strictly conserved GGG-tracts in the 5’ and 3’ long terminal repeat of HIV-1 are critical to control multiple steps of HIV-1 replication to prevent acquisition of unwanted mutations in the region

**DOI:** 10.64898/2026.04.24.720579

**Authors:** Takeshi Yoshida, Yuho Kasuya, Tetsuro Matano, Takao Masuda, Hiroyuki Yamamoto

## Abstract

Three consecutive deoxyguanosine residues (the GGG-tract) in the U3/R junction of the 5’ long terminal repeat (LTR) are strictly conserved among all HIV-1 subtypes. Each deoxyguanosine within the tract has been reported to function as transcription initiation site for HIV-1 RNA. Furthermore, RNAs whose transcription initiates from the third deoxyguanosine in the tract (1G form RNAs) are predominant in virus particles and serve as the primary templates for reverse-transcription. In this study, we generated mutant HIV-1s by replacing the tracts in both the 5’ and 3’ LTR with other nucleotides to elucidate their functional significance. We identified several proviral sequences containing unexpected mutations near the 5’ tract after infection with the mutant, but not the wild-type virus. Five-prime rapid amplification of cDNA end (5’ RACE) analyses of RNAs purified from mutant virus particles revealed multiple RNA variants with 5’ terminal sequences differing from the plasmid used for producing the particles. Some of the unexpected proviral sequences likely arise directly from these variants during reverse-transcription. We also found that replacing all three nucleotides in the 3’ tract with deoxyadenosines decreased the proportion of the 1G form RNAs in particles to 32.6%. Nevertheless, up to 88% of provirus was likely generated with the 1G form RNAs, although they were no longer absolutely predominant in particles. Our results demonstrate that the GGG-tracts in the 5’ and 3’ LTR are conserved to maintain the integrity of reverse-transcription for LTR sequence generation by controlling multiple steps of HIV-1 replication, including transcription, RNA packaging and reverse-transcription.

**Importance:** HIV-1 is highly mutable, yet certain conserved sequences remain consistent across most strains. These sequences are thought to be maintained because mutations in these regions typically impair viral fitness, leading to the elimination of such variants through natural selections. This study suggests that HIV-1 has a unique mechanism to autonomously prevent acquisition of mutations in certain regions. Specifically, the GGG-tracts in the 5’ and 3’ LTR ensure accurate transcription of HIV-1 RNAs, selective packaging of the 1G form RNAs, and preferential use of these 1G forms as a template for reverse-transcription. GGG preservation minimizes unwanted mutations in this region, allowing progeny virus to inherit intact LTR sequences. Interestingly, the preferential usage of 1G form RNAs as template for reverse-transcription is not due to their predominance in virus particles. This suggests that genome packaging independently contributes to reverse-transcription as preparatory stage by concentrating the most suitable RNAs for reverse-transcription in particles.

## Introduction

The ability of HIV-1 to rapidly mutate provides a powerful driving force for escaping from various pressures including human immunity, cellular restriction factors and even antiretroviral drugs (1). However, what actually introduces the mutation remains to be determined, as described previously (2). Although a casual reading of HIV-1 literature would suggest that mutations in HIV-1 genome are introduced by errors made by reverse-transcriptase, the data that address directly to the question are quite limited (2). In theory, there are basically three ways in which HIV-1 genome could acquire mutations, namely, RNA transcription by human DNA-dependent RNA polymerase (RNA Pol II), reverse-transcription by HIV-1 reverse-transcriptase and provirus replication by human DNA polymerase during cell division of infected cells (3). However, it would be reasonable to surmise that there are more opportunities for mutations to be introduced during RNA transcription and reverse-transcription than those during cell division because there might be limited opportunities for most infected cells to divide before they die (2). Furthermore, human DNA polymerase but not human RNA Pol II nor HIV-1 reverse-transcriptase has proofreading function which significantly reduces opportunity of mutations introduced during enzyme-dependent polymerization of nucleotides. In addition to mutations by these enzymes, cellular factors, in particular, the APOBEC proteins are also known to be responsible for accumulating mutations in HIV-1 genome (4).

In transcription of HIV-1 RNA, heterogeneous transcriptional initiation site usage was recently reported as a new mechanism for diversification of HIV-1 RNA functions (5). This mechanism is conserved among most HIV-1 strains (5, 6) and the majority of simian immunodeficiency viruses (7). In the paradigm, fate of the RNAs is primarily determined depending on transcriptional initiation sites among three consecutive deoxyguanosine residues (GGG-tract) downstream of the TATA-box in the 5’ long terminal repeat (LTR) (5, 8). HIV-1 RNA transcription usually starts from the first deoxyguanosine of the GGG-tract but sometimes from the second or third deoxyguanosine (5). However, RNAs beginning with one guanosine (1G form RNAs), whose transcription initiates from the third deoxyguanosine of the tract are predominant in HIV-1 particles (5). The selective packaging of the 1G form RNAs was also observed in virus particles in blood from people living with HIV-1 (9). Interestingly, the secondary and tertiary structure of the 5’ leader sequences in the 1G form RNAs are different with those of the other forms and determine the fate of the RNAs to be dimerized and incorporated into virus particles (10–13).

Our recent study demonstrated that the 1G form RNAs serve as primary templates for reverse-transcription to generate the provirus (14). In the study, we also unexpectedly found that HIV-1 reverse-transcriptase has the ability to overcome mismatches between template RNA and primer DNA and extends the mismatched 3’ termini by incorporation and polymerization of the next complementary nucleotide, in keeping with early reports (15, 16). In fact, after infection with an HIV-1 mutant in which the GGG-tract in the U3/R junction of the 3’ LTR were replaced with TTT, proviral sequences TTG was found to compose 94% of proviral sequences for the region of the GGG-tract in the 5’ LTR (14). Proviral sequence TTG is likely generated with minus-strand strong-stop cDNA (−sscDNA) made with the 1G form RNA combined with the ability of HIV-1 reverse-transcriptase to overcome mismatches (14).

The GGG-tracts in the U3/R junction of the 5’ LTR are strictly conserved among most HIV-1s regardless of subtypes (http://www.hiv.lanl.gov, HIV Sequence Compendium 2023). Conserved sequences within the HIV-1 genome are generally regarded as important regions for surviving strategies of HIV-1 and having been maintained. Conversely, viruses acquiring deleterious mutations to HIV-1 fitness in these regions have likely been eliminated during the selection process. As described previously, each deoxyguanosine residue of the GGG-tract functions as transcription initiation site but the reason why these sequences are strictly conserved among almost all HIV-1 strains are not yet elucidated.

In this study, we found that HIV-1 likely possesses an intrinsic mechanism to avoid acquiring unwanted mutations in specific regions during reverse-transcription even using the low-fidelity reverse-transcriptase. Specifically, the GGG-tract in the 5’ and 3’ LTR were found to be crucial for HIV-1 to make progeny viruses inherit intact sequences of LTR. For achieving the purpose, HIV-1 would use the most suitable RNAs as a template for reverse-transcription, the 1G form RNAs, which are predominantly incorporated into virus particles. Importantly, the 1G form RNAs were primarily used as a template for reverse-transcription but not due to their predominance in particles, suggesting that there are at least two independent selection steps of the 1G form RNAs. Thus, we conclude that the GGG-tracts in the U3/R junction of the 5’ and 3’ LTR are conserved for keeping LTR sequences intact during reverse-transcription by controlling transcription, RNA packaging and reverse-transcription of HIV-1.

## Results

### The GGG-tracts in the U3/R junction of the 5’ LTR and 3’ LTR are not an absolute requisite for attainment of single-round HIV-1 infection

As described in the introduction, each deoxyguanosine residue of the GGG-tract in the U3/R junction of the 5’ LTR functions as transcription initiation site (Fig. 1A). As shown in Figure 1B, the GGG-tract in the 5’ LTR is strictly conserved among almost all HIV-1 strains (http://www.hiv.lanl.gov, HIV Sequence Compendium 2023), but the reason is not yet elucidated. To assess whether the tracts in both the 5’ LTR and 3’ LTR are an absolute requisite for virus replication, we first generated three mutants of single-round HIV clone DNA pNL4-3EGFP*ΔenvΔnef* (17) by replacing the GGG-tracts in both the 5’ and 3’ LTR with deoxyadenosines (the AAA-AAA mutant), deoxycytidines (the CCC-CCC mutant) or deoxythymidines (the TTT-TTT mutant) (Fig. 1C). We transfected 293T cells with each mutant along with a plasmid expressing VSV G protein (pMISSION-VSV-G) to produce the VSV-G-pseudotyped virus. After exposure of T-cell line MT-4 cells to the pseudotyped virus, we assessed whether cells were infected or not by evaluating EGFP expression of MT-4 cells with flow cytometry. Although numbers of infected cells caused with 1 mL of supernatant containing each virus are different (Fig. 1D), these plasmids could produce infectious virus, suggesting that the GGG-tracts in the U3/R junction of the 5’ and 3’ LTR are critical for efficient production of infectious virus but not absolutely essential for HIV-1 production nor infection.

**FIG 1:**
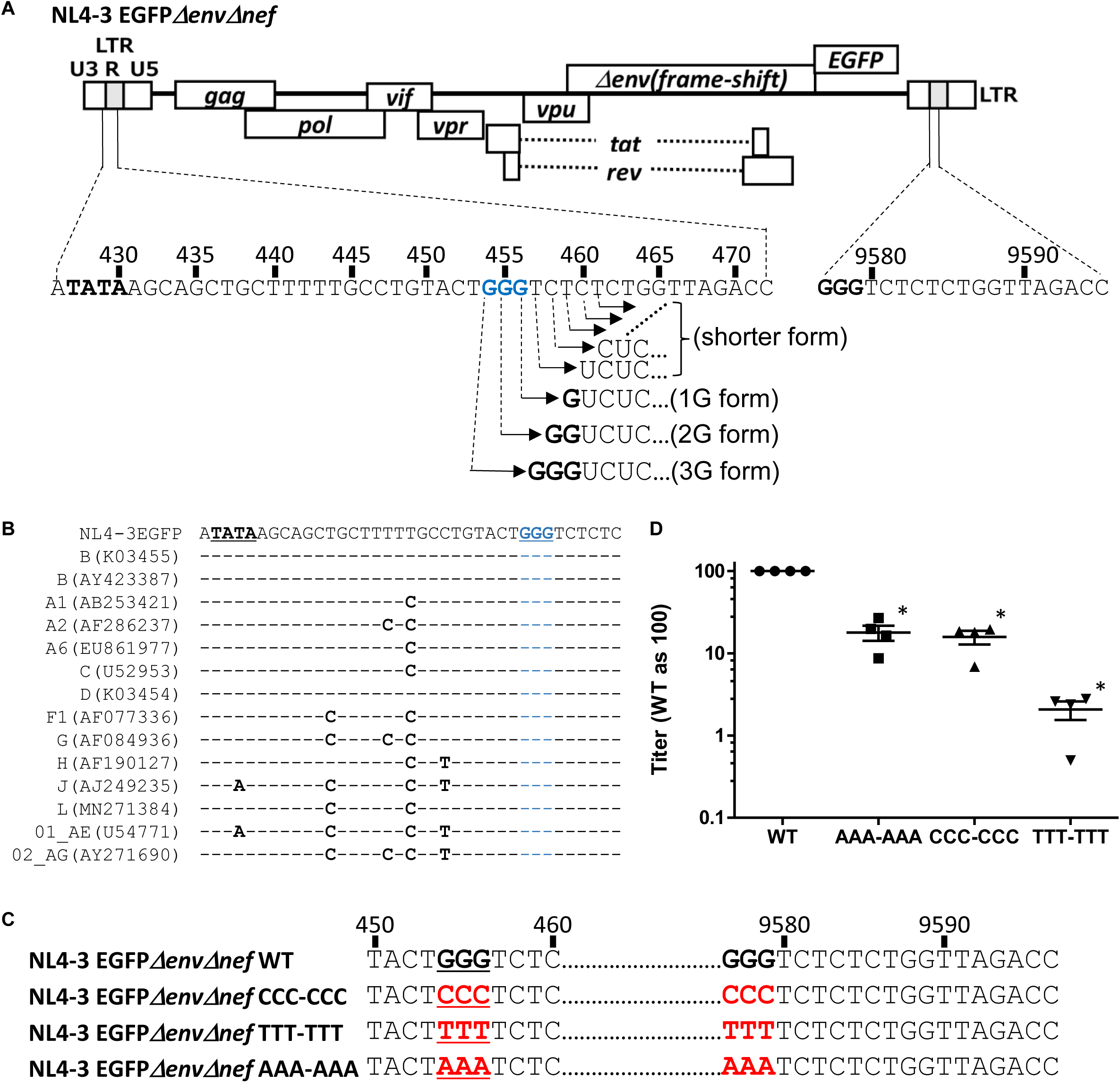
The GGG-tracts in the U3/R junction of both the 5’ and 3’ LTR are critical for efficient production of infectious virus but not absolutely requisite for HIV-1 infection. (A) A schematic represents heterogeneous transcriptional initiation site usage by HIV-1. The strictly conserved GGG-tract is located in the U3/R junction of the 5’ and 3’ LTR of HIV-1 clone DNA (pNL4-3EGFP*ΔenvΔnef*). The 5’ leader sequences of HIV-1 RNA whose transcription initiates from the 1st, 2nd and 3rd deoxyguanosine in the tract are also shown (the 3G, 2G and 1G form, respectively). RNAs whose transcription initiates from nucleotides downstream of the tract are defined as the shorter form RNAs. The TATA-box and the GGG-tracts are highlighted by black and blue bold characters, respectively. (B) Nucleotide sequences between the TATA-box and the GGG-tract in the 5’ LTR of the HIV-1 references from the Los Alamos HIV databases (http://www.hiv.lanl.gov, HIV Sequence Compendium 2023) are shown. Subtypes and accession numbers are shown. Dashes indicate nucleotide identity. The TATA-box and the GGG-tracts are highlighted as black and blue bold characters, respectively. (C) Nucleotide sequences of the CCC-CCC, TTT-TTT and AAA-AAA mutants of NL4-3EGFP*ΔenvΔnef* are shown. The mutated nucleotides are highlighted as red bold characters. The 5’ tract is highlighted with underlines. (D) Numbers of EGFP-positive cells produced with 1 mL of supernatant containing the VSV-G-pseudotyped NL4-3EGFP*ΔenvΔnef* wild-type (WT), AAA-AAA, CCC-CCC, or TTT-TTT mutant virus were evaluated. The numbers for WT were arbitrarily set as 100. Results from four independent experiments are shown. Asterisks code for statistical significance compared to WT: *, P<0.01.

### Unexpected mutations in proviral sequences arise after infection with the GGG-tract double-mutated viruses, but not the wild-type virus

To assess whether provirus can be generated properly by three mutant viruses, we analyzed proviral sequences as done for our preceding study (14). Interestingly, we identified several proviral sequences containing unexpected mutations in the region of the 5’ tract after infection with the CCC-CCC virus. In 9% of clones analyzed, proviral sequences of the region for the 5’ tract were C**G**C (Fig. 2A, 2nd row; an unexpected mutation is highlighted as a bold character) while the other proviruses were CCC (no mutation; Fig. 2A, 1st row). After infection with the TTT-TTT virus, another unexpected mutation TT**C** was also observed (Fig. 2B, 2nd row). Interestingly, some unexpected mutations in provirus were also found in the proximal upstream of the region after infection with the TTT-TTT virus (A**T**T<u>TTT</u>TC: Fig. 2B, 4th row; the region for the 5’ tract is highlighted with an underline) or the AAA-AAA virus (AC**A**<u>AAA</u> and A**A**T<u>AAA</u>: Fig. 2C, 2nd and 3rd row, respectively) and also in the downstream of the tract after infection with the TTT-TTT virus (ACT<u>TTT</u>**G**C: Fig. 2B, 3rd row). In contrast, we did not find any unexpected mutation in proviral sequences after infection with the wild-type virus (the GGG-GGG virus; Fig. 2D).

**FIG 2:**
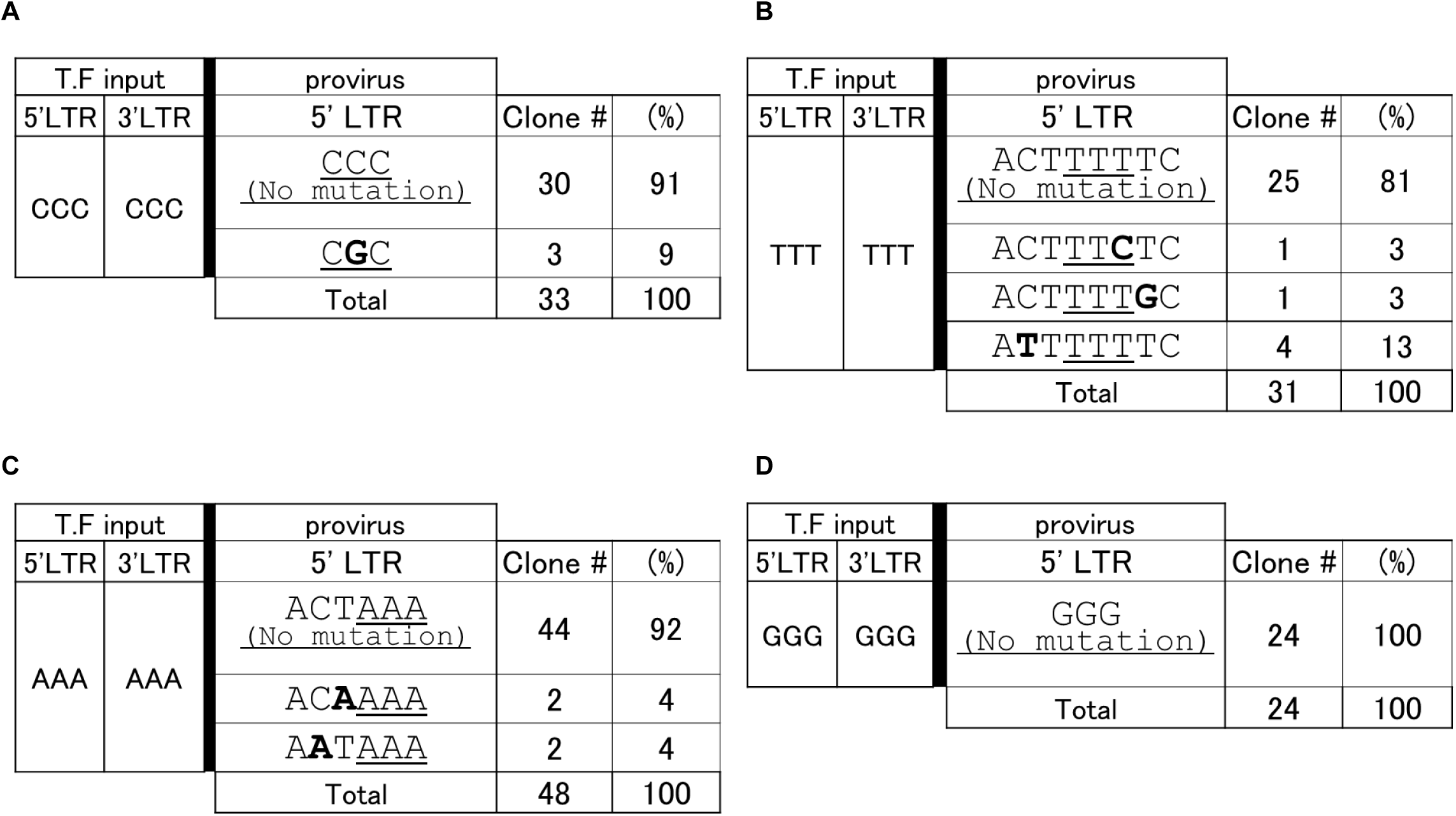
Unexpected mutations in proviral sequences were observed after infection with the GGG-tract double-mutated viruses. (A-D) Sequencing analyses of provirus after infection with the VSV-G-pseudotyped NL4-3EGFP*ΔenvΔnef* CCC-CCC mutant (A; n=33 clones), the TTT-TTT mutant (B; n=31 clones), the AAA-AAA mutant (C; n=48 clones) or the wild-type virus (the GGG-GGG virus; D; n=24 clones) are shown. Unexpected nucleotides and the region of the tract are highlighted with bold letters and underlines, respectively. T. F. input: a plasmid for transfection to express HIV-1 transcripts (pNL4-3EGFP*ΔenvΔnef* WT or mutant plasmid).

### Some unexpected sequences preexist in RNAs incorporated into the mutant virus particles

To understand why some proviral sequences acquired unexpected mutations after infection with mutant virus, a central point to clarify whether these mutations were introduced before or after reverse-transcription. To assess whether RNAs incorporated into virus particles have already acquired mutations, we evaluated the 5’-terminal sequences of genomic RNA incorporated into virus particles by 5’ rapid amplification of cDNA end (5’ RACE) analyses (Fig. 3A). As expected, we found the 1A, 3A and shorter form RNAs in particles of the AAA-AAA mutant virus (Fig. 3B, above the bold black bar). In the same sample, we also found small numbers of unexpected forms of RNA including the 4A (**A**AAA), 1C (**C**), 1G (**G**), **G**AAA, and **G**AA form RNAs (Fig. 3B, below the bold black bar) while in particles of the wild-type virus (GGG-GGG virus), no RNA forms other than the 1G, 2G, 3G and shorter form were detected in our previous study (14). In each of these unexpected forms, one nucleotide differs from DNA sequences of the parental plasmid of the AAA-AAA mutant (TACT<u>AAA</u> in Fig. 1C, 4th row; different nucleotides are highlighted as bold characters).

**FIG 3:**
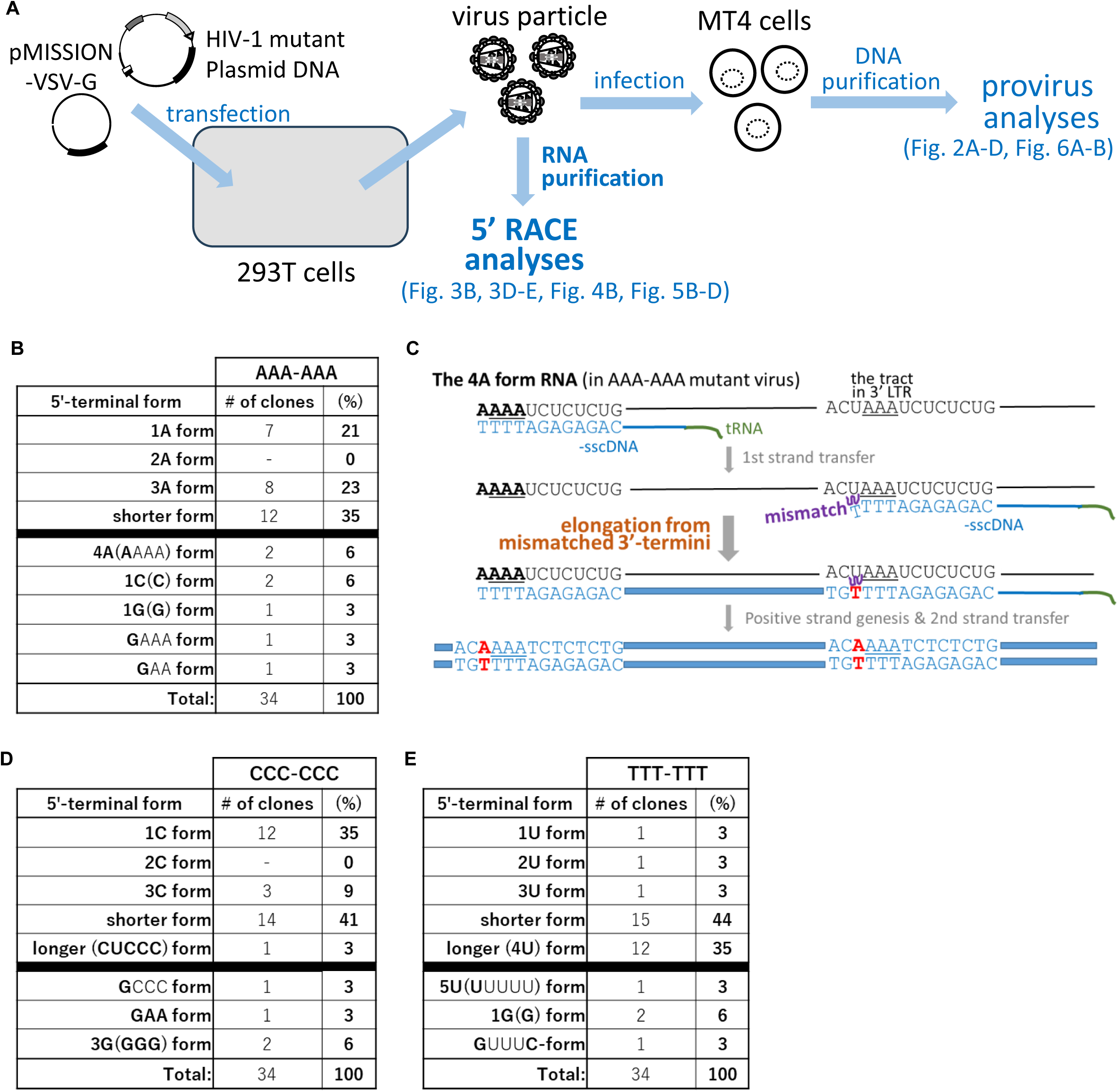
Unexpected sequences were also found in RNAs incorporated into the mutant virus particles. (A) A schematic represents 5’ RACE analyses and provirus analyses. 293T cells were transfected with an HIV-1 clone plasmid (pNL4-3EGFP*ΔenvΔnef* WT or mutant plasmid) and pMISSION-VSV-G to produce the VSV-G-pseudotyped HIV-1. Genomic RNAs in the particles were purified and subsequently subjected to 5’ RACE. For provirus analyses, MT-4 cells were exposed with supernatant containing the VSV-G-pseudotyped virus. Genomic DNA containing HIV-1 provirus were purified from MT-4 cells. (B) Results of 5’ RACE analyses are shown. 5’RACE of purified RNA from the VSV-G-pseudotyped virus particles were done for the AAA-AAA mutant virus (n=34 clones). Nucleotides different from the input plasmid for producing particles (Fig. 1A and 1C) are highlighted as bold characters. Hyphens are used for forms not observed in the analyses. (C) HIV-1 reverse-transcriptase has been reported to have the ability to overcome mismatched 3’ termini between a template RNA and minus-strand strong-stop cDNA (-sscDNA). A schematic based on the knowledge represents predicted reverse-transcription processes to generate unexpected proviral sequences (AC**A**<u>AAA;</u> the region of the tract is highlighted with an underline) with the 4A form template RNA. Genomic RNAs, DNAs, acquired mutations and tRNAs are drawn in black, blue, red and green, respectively. (D-E) Results of 5’ RACE analyses are shown as described in Figure 3B. The analyses of purified RNA from the particles were done for the CCC-CCC (C; n=34 clones) or the TTT-TTT mutant virus (D; n=34 clones).

In our previous study (14), we unexpectedly found that HIV-1 reverse-transcriptase overcame a mismatch between a template RNA and 3’ termini of primer DNA by polymerizing the next complementary nucleotide DNA over the mismatch, recapitulating early reports (15, 16). As shown in Figure 3C, reverse-transcription with the 4A form template RNAs (Fig. 3B, 5th row) would likely generate one of unexpected proviral sequences, AC**A**<u>AAA</u> (Fig. 2C, 2nd row). On the other hand, the **A**TAAA form RNA which would be a template for the other unexpected proviral sequences A**A**T<u>AAA</u> (Fig. 2C, 3rd row) was not found in the AAA-AAA mutant virus particles. Similarly, in particles of the CCC-CCC mutant virus (Fig. 3D), there are unexpected forms including the **G**CCC, **GAA** and 3G (**GGG**) form RNAs but not the **G**C form RNA which likely produce the unexpected proviral sequences <u>C**G**C</u> (Fig. 2A, 2nd row). In the TTT-TTT mutant virus particles, unexpected forms of RNA including the 5U (**U**U<u>UUU</u>; the region for the 5’ tract is highlighted with an underline), 1G (**G**) and **G**UUU**C** form RNAs (Fig. 3E, below the bold black bar) in addition to the 1U, 2U, 3U, shorter and longer (4U) form RNAs were found (Fig. 3E, above the bold black bar). Although the 5U form RNAs are predicted to produce one of the unexpected proviral sequences (A**T**T<u>TTT</u>TC in Fig. 2B, 4th row), RNAs for the other unexpected proviral sequences (ACT<u>TT**C**</u>TC and ACT<u>TTT</u>**G**C) were not observed in the assay. From the series of 5’ RACE assay, a number of RNAs with unexpected sequences in the 5’-termini were found and some can be a template RNA for reverse-transcription to generate some unexpected proviral sequences shown in Figure 2A-2C. Thus, HIV-1 could acquire some mutations in its genome RNAs likely preceding reverse-transcription when HIV-1 genome lacks the GGG-tracts in both the 5’ and 3’ LTR.

### The GGG-tracts in the U3/R junction of the 3’ LTR are critical for RNA selection during RNA packaging into virus particles

Interestingly, trend of RNAs incorporated into the AAA-AAA mutant virus particles were found to be different from those in the AAA-GGG mutant virus particles in our previous study [the NL4-3EGFP *Δenv Δnef* AAA mutant in (14), in which the 5’ GGG-tract was replaced with deoxyadenosines]. There are the 3A form RNAs in the AAA-AAA mutant but not in the AAA-GGG mutant virus particles [23% in Figure 3B and 0% in (14), raw data is not shown here, respectively]. An exact opposite phenomenon was observed for the 2A form RNAs; the 2A form RNAs were not found in the AAA-AAA mutant virus particles (Fig. 3B) but compose 11% of the total RNA in the AAA-GGG mutant virus in our previous study [(14), raw data is not shown here]. Since similar phenomena were also observed for the TTT-TTT and TTT-GGG [the NL4-3EGFP *Δenv Δnef* TTT mutant in (14)] mutant virus, we hypothesized whether the 3’ tract (Fig. 1A, right side) might be critical for RNA selection during RNA packaging into virus particles.

To assess the possibility, we generated the indicated three mutant plasmids, the CCC-AAA, CCC-GGG, CCC-TTT (Fig. 4A). After production of mutant virus particles with them, we carried out 5’ RACE analyses as described in Figure 3 (a schematic is shown in Fig. 3A). As shown below the bold black bars in Figure 4B, multiple unexpected forms of RNA were found. Interestingly, each unexpected form of RNA has one or some nucleotide(s) which is/are different from the input plasmid DNA used for producing virus particles (Fig. 4A, 450-460). The overall incorporation profile of RNAs in the CCC-CCC mutant (Fig. 3D) is similar to that of the CCC-TTT mutant, but distinct from those of the CCC-AAA and CCC-GGG mutants when we focus on the expected RNAs from accurate transcription (Fig. 4B, above the bold black bars). Although the 1C and shorter forms are rich in the CCC-CCC and CCC-TTT mutant virus particles, exact occupied proportion by the 1C and shorter form is different in the CCC-CCC (Fig. 3D; 35% and 41%, respectively) and the CCC-TTT (Fig. 4B, right panels; 60.5% and 27%, respectively) mutant. These results suggest that selection of incorporated RNA into virus particles is affected by the 3 nucleotides in the U3/R junction of the 3’ LTR.

**FIG 4:**
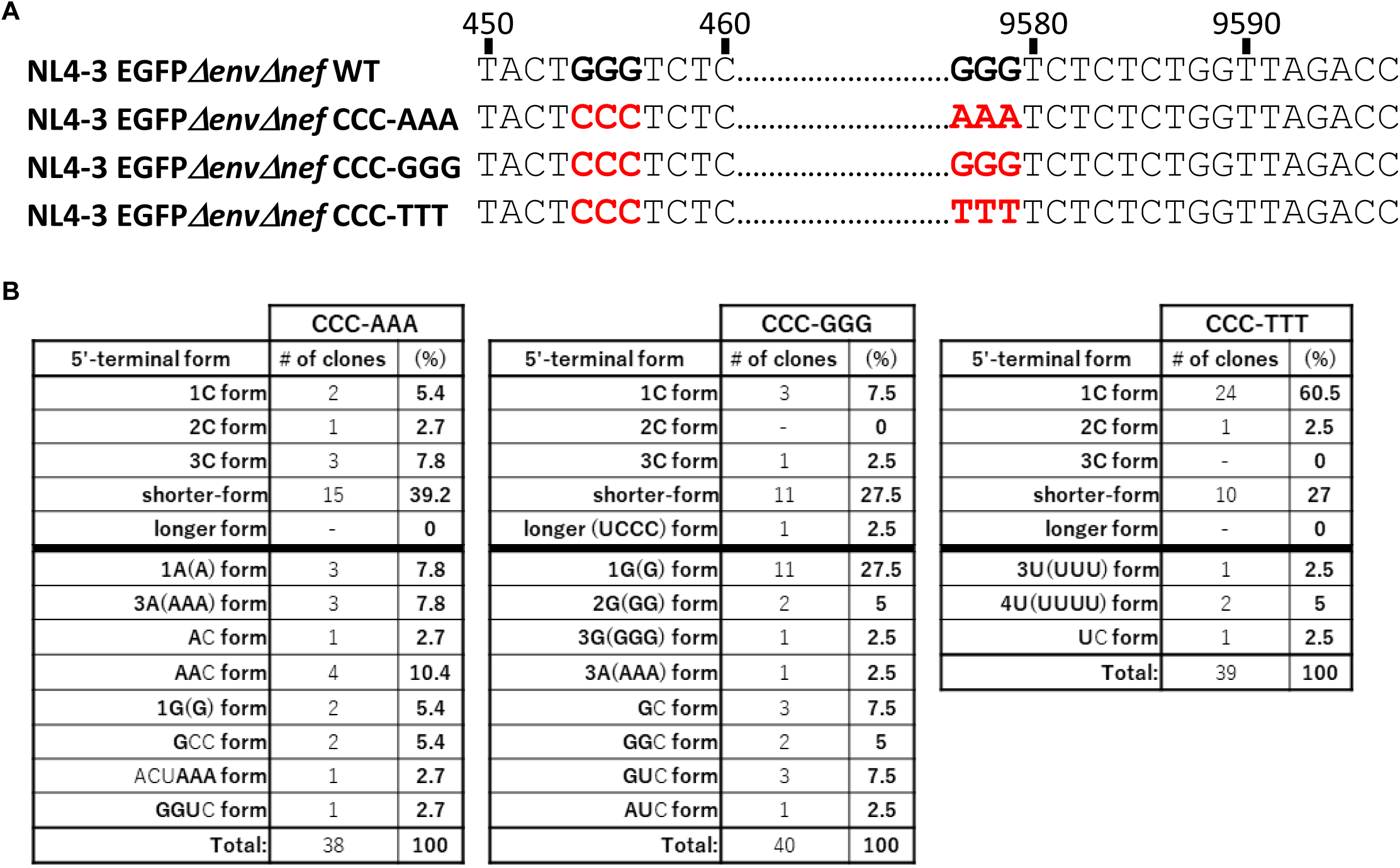
The tract in the U3/R junction of the 3’ LTR is critical for RNA selection during RNA packaging into the virus particles. (A) Nucleotide sequences of the CCC-AAA, CCC-GGG and CCC-TTT mutants of NL4-3EGFP*ΔenvΔnef* are shown as described in Figure 1C. (B) Results of 5’ RACE analyses are shown as described in Figure 3B. 5’ RACE analyses of purified RNA from the particles were done as described in Figure 3A for the CCC-AAA (left panels; n=38 clones), the CCC-GGG (center panels; n=40 clones) and the CCC-TTT mutant virus (right panels; n=39 clones).

Although the 1G form RNAs of HIV-1 are known to account for approximately 75-85% in genomic RNAs in the wild-type (the GGG-GGG) virus particles (5, 7, 14, 18, 19), we next questioned that mutations in the 3’ tract might change this strict regulation. To assess the possibility, we generated another set of three mutants of the 3’ tract, the GGG-AAA, GGG-CCC, and GGG-TTT (Fig. 5A), and carried out 5’ RACE analyses as described in Figure 3 (a schematic is shown in Fig. 3A). As shown in Figure 5B, 5C and 5D, the 1G form RNAs accounted for 32.6%, 60% and 50% in particles of the GGG-AAA, GGG-CCC and GGG-TTT mutant virus, respectively. From these results, we concluded that the 3’ tract are important for the strict regulation that the 1G form RNAs occupy 75-85% of RNA in particles of the wild-type virus, suggesting that it would be thus critical for RNA packaging into virus particles.

**FIG 5:**
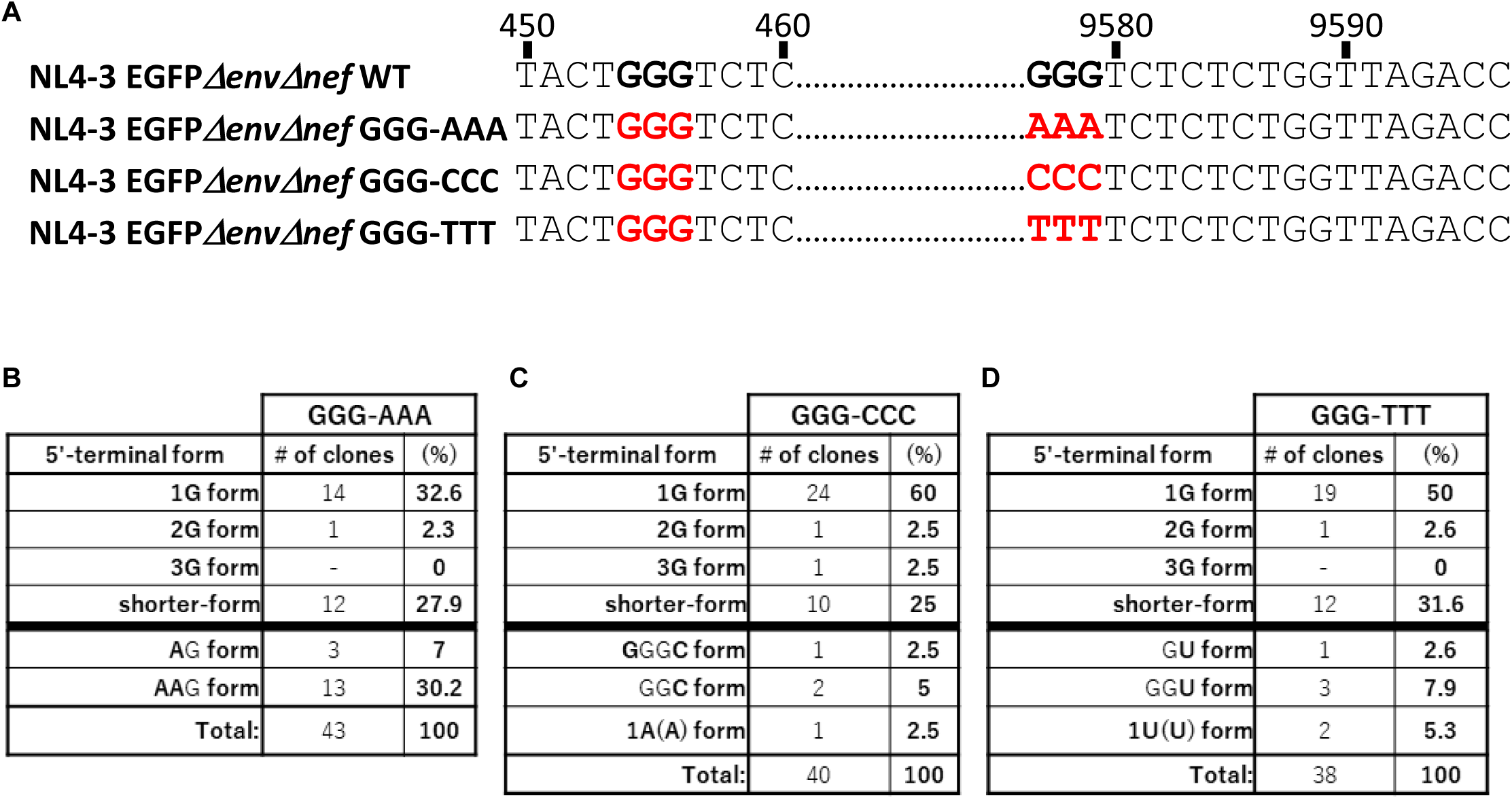
The GGG-tract in the U3/R junction of the 3’ LTR are critical for strict selection of the 1G form RNAs during RNA packaging into virus particles. (A) Nucleotide sequences of the GGG-AAA, GGG-CCC and GGG-TTT mutants of NL4-3EGFP*ΔenvΔnef* are shown as described in Figure 1C. (B-D) Results of 5’ RACE analyses are shown as described in Figure 3B. 5’ RACE analyses of purified RNA from the particles were done as described in Figure 3A for the GGG-AAA (B; n=43 clones), the GGG-CCC (C; n=40 clones) and the GGG-TTT mutant virus (D; n=38 clones).

### Predominance of proviral sequences predicted to be generated with the 1G form RNAs

Since the 1G form RNAs accounted for only 32.6% of RNAs in particles of the GGG-AAA mutant virus, we asked whether the 1G form RNAs might not serve as primary templates for reverse-transcription under such condition. To assess the possibility, we analyzed provirus sequences after infection with the GGG-AAA and also the GGG-CCC mutant virus as done for Figure 2 (a schematic is shown in Fig. 3A). From the results, proviral sequences AAA, AAG and AGG in the region of the 5’ tract were found to compose 6%, 88% and 6% of all provirus after infection with the GGG-AAA mutant virus, respectively (Fig. 6A). Proviral sequences AAA, AAG and AGG are predicted to be produced with the shorter, 1G, and 2G form RNAs, respectively (the reverse-transcription processes are shown in Fig. 3C). These results indicate that the 1G form RNAs would be primarily used as a template for reverse-transcription to generate proviral DNA even though they composed only 32.6% of the total RNA in particles. In the case of the GGG-CCC mutant virus, proviral sequences CCG were found to account for 75% (Fig. 6B, 3rd row). From these analyses, the 1G form RNAs are found to serve as primary templates for reverse-transcription to generate provirus, suggesting that the preferential usage of the 1G form RNAs as a template is not due to their predominance in particles. Of note, proviral sequences with an unexpected mutation, <u>CCC</u>**G**C was also found after infection with the GGG-CCC mutant virus (Fig. 6B, 5th row).

**FIG 6:**
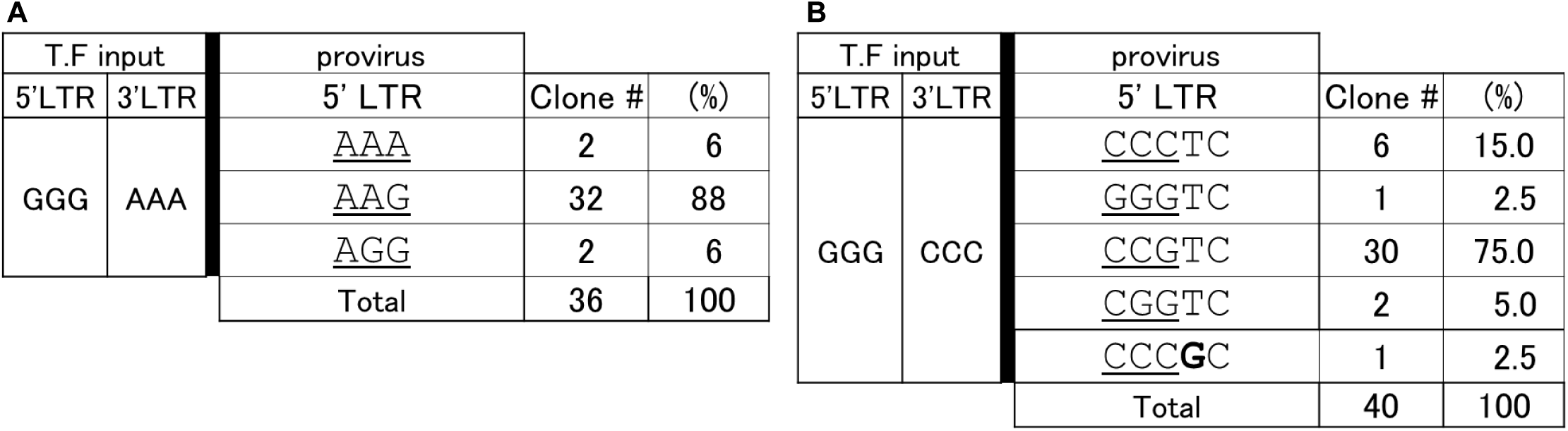
Proviral sequences predicted to be generated with the 1G form RNAs were dominantly found. (A-B) Results of sequencing analyses are shown as described in Figure 2A. The analyses of proviral sequences after infection with the VSV-G-pseudotyped NL4-3EGFP*ΔenvΔnef* GGG-AAA (A; n=36 clones) or the GGG-CCC (B; n=40 clones) mutant virus were done as described in Figure 3A.

## Discussion

From Figure 1D, we found that plasmids for mutant HIV-1 clones by replacing the GGG-tracts in both the 5’ and 3’ LTR with deoxyadenosine, deoxycytidines or deoxythymidine (the AAA-AAA, CCC-CCC, or TTT-TTT mutant) produced infectious virus, although number of infected cells produced with 1 mL of supernatant containing each mutant virus were approximately one fifth, one sixth, or one twentieth of those with the wild-type virus, respectively. It might be reasonable because replacement of the GGG-tract in 5’LTR with AAA or TTT was found not to eliminate viral infectivity in our previous study (14). Yet independent of this, it is unexpected that a subpopulation of proviral sequences acquired mutations in region of the 5’ tract and in its upstream or downstream upon next-round infection with each mutant virus (Fig. 2A-2C). Since no mutation was found in proviral sequences after infection with the wild-type virus (the GGG-GGG virus; Fig. 2D), the GGG-tracts in the U3/R junction of the 5’ and 3’ LTR appear to be critical for making progeny viruses inherit intact genomic information especially sequences of LTR. Results of this functional assay matches the fact that the GGG-tracts are strictly conserved among HIV-1 isolates (Fig. 1B). As shown in Figure 3C, we reported the mechanism of HIV-1 acquiring mutations in the region of the 5’ tract when there are differences between the tracts of the 5’ LTR and 3’ LTR such as the GGG-TTT mutant in previous study (14), but there is no difference in the current mutants including the CCC-CCC, TTT-TTT or AAA-AAA virus (Fig. 1C). If all processes including transcription, RNA packaging and reverse-transcription had been done properly, provirus would not have acquired mutations.

From our 5’ RACE analyses, we found the 4A (**A**AAA) form RNAs in particles of the AAA-AAA mutant virus (Fig. 3B, 5th row) and the 5U (**U**UUUU) form RNA in particles of the TTT-TTT mutant virus (Fig. 3E, 6th row) which are predicted to generate unexpected proviruses such as AC**A**<u>AAA</u> (Fig. 2C, 2nd row) and A**T**T<u>TTT</u>TC (Fig. 2B, 4th row) after reverse-transcription, respectively (the processes are shown in Fig. 3C). If transcription of HIV-1 RNA was done properly, there would not be the 4A (**A**AAA) form RNAs in the AAA-AAA mutant particles nor the 5U (**U**UUUU) form RNAs in the TTT-TTT mutant particles because nucleotides highlighted as bold characters in the RNAs are missing in the sequences between the TATA-box and the 5’ GGG-tract in the parental HIV-1 clone plasmid DNA (Fig. 1C). These results suggest that some unexpected mutations in proviral sequences were most likely produced by unexpected forms of viral genomic RNA. Although we currently do not know the reason why these unexpected form RNAs were found, we can raise two possibilities including errors during RNA transcription and a post-transcriptional addition of one ribonucleotide to the 5’-terminus of genomic RNA. Specifically, the 4A (**A**AAA) or 5U (**U**UUUU) form RNAs are possibly caused after transfer of an adenine or uracil to the 5’-terminus of the 3A form or 4U form RNAs, respectively (one base upstream of the 5’ TTT-tract in the TTT-TTT mutant is also T in Figure 1C). On the other hand, we could not find possible template RNAs for the other unexpected proviral sequences including <u>C**G**C</u> (Fig. 2A, 2nd row; the **GC** form RNAs), A**A**T<u>AAA</u> (Fig. 2B, 3rd row; the **A**TAAA form RNAs), and both ACT<u>TT**C**</u>TC and ACT<u>TTT</u>**G**C [Fig. 2C, 2nd and 3rd row; the 1C (**C**) and unexpected shorter (**G**CUC) form RNAs] after infection with the CCC-CCC, AAA-AAA, and TTT-TTT mutant virus, respectively. Therefore, HIV-1 can acquire mutations due to errors during RNA transcription or post-transcriptional modification, but other mechanisms may be also involved.

Interestingly, two previous studies also showed one of similar phenomena of mutations in the tract after replication of an HIV-1 mutant, in which the GGG-tract was replaced with TTG (20) or TCG (21). Since more than half of proviral sequences of the 5’ tract was changed to TGG from TTG after serial passaging, authors proposed the hypothesis that guanosine cap of the 1G form RNAs might function as a template for reverse-transcription, non-encoded deoxycytidine might be added to the 3’ terminus of minus-strand strong-stop cDNA (−sscDNA), and finally proviral sequences TGG at the 5’ tract might be generated after revers-transcription (20). Another study also assessed this possibility through *in vitro* reverse-transcription and concluded that ^m7^G of the cap on HIV-1 RNA might not be reverse-transcribed in the system (22). In our study, we also found some mutations replaced with deoxyguanosine residue such as <u>C**G**C</u> after infection with the CCC-CCC mutant virus (Fig. 2A, 2nd row), ACT<u>TTT</u>**G**C after infection with the TTT-TTT mutant virus (Fig. 2B, 3rd row), and <u>CCC</u>**G**C after infection with the GGG-CCC mutant virus (Fig. 6B, 5th row). Although many types of RNA variants with 5’ terminal sequences differing from the parental plasmid were found, we could not identify the source RNAs for the unexpected proviral sequences shown here. Interestingly, 1 nucleotide downstream of the unexpectedly appearing deoxyguanosine residue is always deoxycytidine in these three cases of this study (<u>C**G**C</u>, ACT<u>TTT</u>**G**C, and <u>CCC</u>**G**C).

HIV-1 is known to be highly mutative and evolve so rapidly to fit *in vivo* environment in persistent infection. Although evolution consists of acquiring mutations and selections, it is difficult to define mutations introduced during HIV-1 replication in the absence of selections because there is selection pressure even in its replication in cell lines. Since we employed single-round infection with a VSV-G pseudotyped HIV-1, we were able to observe not only advantageous mutations but also synonymous and disadvantageous mutations. We purified RNAs not only from infectious virus particles but from all virus particles including non-infectious particles. We also extracted proviral DNAs not only from cells expressing viral proteins and/or producing infectious progeny virus but all infected cells. It might be possible that this approach counterintuitively enabled us to uncover the property avoiding mutations by the strictly conserved GGG-tracts.

As shown in Figures 5B-5D, replacement of the GGG-tract in the 3’ LTR even with an intact 5’ GGG-tract considerably weakened the bias to select the 1G form RNAs during RNA packaging. In fact, the occupied proportion by the 1G form RNAs are 32.6%, 60% and 50% in particles of the GGG-AAA, GGG-CCC and GGG-TTT mutant virus (Fig. 5B, 5C and 5D, respectively). Thus, we conclude that not only the GGG-tract in the 5’ LTR but also that in the 3’ LTR are critical for RNA selection during RNA packaging. Although it is difficult to predict how the 3’ tract contributes to RNA packaging into virus particles, a previous report by Brown et al. (10) can be a hint. Since the secondary and tertiary structure of the HIV-1 5’ leader sequences was reported to control RNA selection on the RNA packaging (10), it is possible to hypothesize that the 3’ tract could affect the structure directly or indirectly.

In Figure 6, proviral sequences AAG and CCG which are predicted to be generated with the 1G form RNAs were found to account for 88% and 75% after infection with the GGG-AAA and GGG-CCC mutant virus, respectively. In the case of the GGG-AAA mutant virus, the 1G form RNAs composed only 32.6% of the total RNA in virus particles (Fig. 5B, 1st row) while proviral sequences AAG accounted for 88% (Fig. 6A, 2nd row), suggesting that preferential use of the 1G RNAs as a template for reverse-transcription is not necessarily because they are predominant in virus particles. Similar phenomena were also observed in the GGG-CCC mutant virus that the 1G form RNAs composed 60% of the total RNAs in particles (Fig. 5C, 1st row), but proviral sequences CCG accounted for 75% (Fig. 6B, 3rd row). The shorter form RNAs are predicted to be advantageous on reverse-transcription because minus-strand strong-stop cDNA (−sscDNA) generated with it can completely match the 3’ R region of template RNA after 1st strand transfer while -sscDNA generated with the 1G form RNAs caused a mismatch with nucleotides in 3’ R region, AA<u>A</u> or UU<u>U</u> (mismatched nucleotides with -sscDNA generated with the 1G form RNAs are highlighted with underlines). Even in the situation, the 1G form RNAs are the most used RNAs as a template for generating provirus DNA. From these results, there are two independent selections of the 1G form RNAs during replication of the wild-type virus but regulation to select them during reverse-transcription would be stricter than that of RNA packaging. Thus, it is suggested that RNA packaging into particles functions as the preparatory stage of reverse-transcription by independently concentrating the 1G from RNAs in particles. RNA packaging might reasonably have a functional connection with reverse-transcription process especially in the recently proposed paradigm in which most reverse-transcription processes are done within an intact capsid core (23).

Of note, we found the longer form RNAs whose transcription initiated from a nucleotide upstream of the 5’ tract in the CCC-CCC mutant (the CUCCC form; Fig. 3D, 5th row), the TTT-TTT mutant (the 4U form; Fig, 3E, 5th row), and the CCC-GGG mutant (the UCCC form; Fig. 4B, middle panels, 5th row) virus particles. Interestingly, the longer form RNA was not found in the mutants with the intact GGG-tract in the 5’ LTR including the GGG-AAA, GGG-CCC and GGG-TTT mutant (Fig. 5B-5D). It is likely the evidence that the GGG-tract in the 5’ LTR functions for strict determination of transcription initiation site.

Overall, the GGG-tracts in the U3/R junction of the 5’ and 3’ LTR are found to be critical for preventing unwanted mutations in the region and making progeny viruses inherit intact genomic information especially LTR sequences by controlling multiple steps of HIV-1 replication, including transcription, RNA packaging and reverse-transcription. While HIV-1 is known for its high mutation rate, the 5’ and 3’ GGG-tracts are conserved, potentially acting as structural anchors that maintain intact LTR sequences, much like a solid frame supporting a fluid interior.

## Material and Methods

### Cells and transfection

The methods were described previously (14). HEK293T cells were maintained in Dulbecco’s Modified Eagle Medium (DMEM) (Gibco, Waltham, MA) supplemented with 10% heat-inactivated fetal bovine serum (GE Healthcare, Logan, UT) and penicillin-streptomycin (Fujifilm Wako, Osaka, Japan) at 37 °C with 5% CO_2_. MT-4 cells were maintained in RPMI-1640 Medium (Gibco) supplemented with 10% heat-inactivated fetal bovine serum (GE Healthcare) and penicillin-streptomycin (Fujifilm Wako) at 37 °C with 5% CO_2_. HEK293T cells were transfected using polyethylenimine (PEI, PolyScience, Niles, IL).

### Plasmids

pNL4-3EGFP*ΔenvΔnef* WT and the GGG-TTT (named as the 3’LTR TTT mutant in the reference) was described previously (14). The mutants of pNL4-3EGFP*ΔenvΔnef* (the CCC-CCC, TTT-TTT, AAA-AAA, CCC-AAA, CCC-GGG, CCC-TTT, GGG-AAA and GGG-CCC) were constructed by overlap-extension PCR. All mutations were confirmed by sequencing analyses.

### Evaluation of infected cells

The methods are described previously (14). To produce VSV-G pseudotyped HIV-1 (24), approximately 2×10^6^ of 293T cells grown in 6-well plate were transfected with 3 μg of proviral plasmid pNL4-3EGFP*ΔenvΔnef* WT or mutants and 1 μg of pMISSION-VSV-G (Sigma Aldrich, St. Louis, MO). Twenty-four h post-transfection, culture medium was changed once. Forty-eight h post-transfection, the supernatants containing VSV-G-pseudotyped HIV-1 were harvested and filtered through 0.45 μm-pore filters (Merck Millipore, Burlington, MA). Approximately 2×10^6^ of MT-4 cells were exposed with an optimized amount (achieving less than 30% EGFP-positive population) of supernatant containing virus. At 48 h after infection, cells were harvested and analyzed on FACSCalibur flow cytometer and analyzed using BD Cell Quest Pro software (BD Bioscience, San Diego, CA).

### Sequencing of provirus DNA

The methods were described previously (14). The harvested supernatants containing the VSV-G-pseudotyped HIV-1 were treated with RNase-free Recombinant DNase I (Takara Bio, Shiga, Japan) at 37 °C for 20 min. At 37 °C, approximately 2×10^6^ of MT-4 cells were incubated with DNase I-treated supernatant in a 15 ml tube for 60 min and washed with 10 ml of medium three times. Forty-eight h post infection, EGFP expression was confirmed by microscope and genomic DNA was purified using DNeasy Blood & Tissue Kit (Qiagen Inc, Hilden, Germany). The 1st PCR was done with Prime STAR HS (Takara) and primers shown below. seqFW238-NL: 5’-CGGAGGGAGAAGTATTAGTG-3’, seqRV998gag-NL: 5’-GTCTGAAGGGATGGTTGTAG-3’ could amplify sequences including 5’ tract but not those including 3’ tract. The cycling condition included a denaturation step (94°C for 1 min), followed by 35 cycles of denaturation (94°C for 30 s), annealing (52°C for 30 s), and extension (72°C for 50 s). The 1st PCR product was subsequently subjected to the 2nd PCR using primers (seqFW266-NL: 5’-GACAGCCTCCTAGCATTTC-3’, seqRV964gag-NL: 5’-GTCTACAGCCTTCTGATGTC-3’) with the same condition except for annealing (60°C for 30 s). DNA fragments with appropriate size from the 2nd PCR product was cloned into pCR4 using Zero Blunt™ TOPO™ PCR Cloning Kit (Invitrogen, Waltham, MA). Sequences of cloned fragments were analyzed by Eurofins Genomics (Tokyo, Japan). More than 24 clones were analyzed per mutant.

### 5’ RACE Assay

The methods were described previously (14). Approximately 7×10^6^ of 293T cells grown in a 10 cm plate (Corning Inc., Durham, NC) were transfected with 15 μg of the HIV-1 plasmid (pNL4-3EGFP*ΔenvΔnef* mutants) and 6 μg of pMISSION-VSV-G (Sigma Aldrich) and culture medium was changed three times 24 h post-transfection. At 48 h after transfection, supernatants containing VSV-G-pseudotyped virus were harvested, filtered through 0.45 μm-pore of filters (Merck Millipore) and concentrated using PEG-IT (System Biosciences, Mountain View, CA). Viral RNA in virus particles were purified using Isogen (Fujifilm Wako) and purified RNAs were subjected to SMARTer® RACE 5’/3’ Kit (Takara Bio USA Inc., San Jose, CA). As shown in Rawson et al. (7), RT primer: 5’-GGTGGCTCCTTCTGATAATG-3’, and reverse PCR primer: 5’-GATTACGCCAAGCTTTCGTTCTAGCTCCCTGCTTG-3’ were used. Sequences of cloned fragments were analyzed by Eurofins Genomics. More than 34 clones were analyzed per mutant.

### Statistical Analysis

Statistical analyses were performed using GraphPad Prism 6. Data are presented as means with error bars indicating SEM from four independent experiments. One-way analysis of variance (ANOVA) with Dunnett’s multiple comparison test was used for comparison of titer with the wild-type virus (WT; Fig. 1D). (Asterisk code for statistical significance: *, *P*<0.01).

## Acknowledgements

T.Y. thanks Dr. Yuko Yoshida for fruitful discussion. This work was supported by a grant 24fk0410064j0401 to T.Y. by the Japan Agency for Medical Research and Development (AMED) Research Project on AIDS/HIV, and a grant 1224 to T.Y. from Takeda Science Foundation. We also recognize support from Dr. Masako Nishizawa, Dr. Takaya Hayshi, Dr. Koji Sakai, Dr. Shoji Yamaoka, Dr. Shigeyoshi Harada, Dr. Sayuri Seki and Ms. Shioko Kojima.

## Author contributions

TY conceived the study and designed experiments. TY and YK performed experiments. TY, YK, T. Matano, T. Masuda and HY analyzed the data. TY and T. Masuda contributed to reagents and materials. TY wrote the paper. All authors reviewed the manuscript.

